# Sequence diversity and evolution of an iflavirus family associated with ticks

**DOI:** 10.1101/2020.06.02.129551

**Authors:** Romain Daveu, Caroline Hervet, Louane Sigrist, Davide Sassera, Aaron Jex, Karine Labadie, Jean-Marc Aury, Olivier Plantard, Claude Rispe

**Affiliations:** INRAE, Oniris, BIOEPAR, Nantes, France; Department of Biology and Biotechnology “L. Spallanzani”, University of Pavia, Pavia, Italy; Population Health and Immunity Division, Walter and Eliza Hall Institute of Medical Research, Parkville, Victoria, 3052, Australia; Faculty of Veterinary and Agricultural Sciences, The University of Melbourne, Parkville, Victoria, 3010, Australia; Genoscope, Institut de biologie François-Jacob, Commissariat à l’Energie Atomique (CEA), Université Paris-Saclay, Evry, France; Génomique Métabolique, Genoscope, Institut François Jacob, CEA, CNRS, Univ Evry, Université Paris-Saclay, 91057 Evry, France

**Keywords:** Virus, Arthropods, Ixodes ricinus, Transcriptome, RNA-Seq

## Abstract

We studied a family of iflaviruses, a group of RNA viruses frequently found in arthropods, focusing on viruses associated with ticks. Our aim was to bring insight on the evolutionary dynamics of this group of viruses, which may interact with the biology of ticks. We explored systematically *de novo* RNA-Seq assemblies available for species of ticks which allowed to identify nine new genomes of iflaviruses. The phylogeny of virus sequences was not congruent with that of the tick hosts, suggesting recurrent host changes across tick genera along evolution. We identified five different variants with a complete or near-complete genome in *Ixodes ricinus.* These sequences were closely related, which allowed a fine-scale estimation of patterns of substitutions: we detected a strong excess of synonymous mutations suggesting evolution under strong positive selection. ISIV, a sequence found in the ISE6 cell line of *Ixodes scapularis,* was unexpectedly nearidentical with *I. ricinus* variants, suggesting a contamination of this cell line by *I. ricinus* material. Overall, our work constitutes a step in the understanding of the interactions between this family of viruses and ticks.

## 1. Introduction

Ticks are blood-feeding parasites, which makes of them a “hub” for a number of microorganisms (bacterial pathogens and symbionts, viruses, etc.) potentially interacting with ticks and their vertebrate hosts. We will here focus on one of these interactors, RNA viruses, and especially a family of iflaviruses which has recently been shown to represent an occasional partner of ticks. More broadly, RNA viruses have accompanied the evolution of cellular life from its very beginning or even preceded it [1]. Beyond the restricted number of viruses causing diseases to humans or agronomical species, which have concentrated more studies so far, much remains to discover on the impact of viruses on other organisms and on their evolutionary trajectories. The discovery of new RNA viruses has rapidly increased in the recent years, and they have appeared to be far more prevalent and diverse than previously expected, especially in invertebrates, which exhibit a large and diverse “virosphere” [2,3]. High throughput transcriptome sequencing (or RNA-Seq) has proven to be a tool of choice to characterize the genomes of these viruses (see [4], [3], and even in ticks [5], [6], as this technique does not require to target known sequences, while allowing to retrieve contigs that may contain full viral genomes given a sufficient coverage and sufficient prevalence of the virus in the analyzed tissues/individuals. Iflaviruses represent a particular group of RNA viruses: members of Picornavirales, they are non-envelopped and have a single strand positive sense non segmented genome of ~9-11 kb – see [7] for a general description. This family of viruses has been shown to have a privileged association with insects, being detected in a growing number of species of several orders. The symptoms associated with this infection vary from null to developmental anomalies (deformed wing virus), or mortality [7]. In addition, iflaviruses have also been detected in some Chelicerata, especially in parasitic mites (e.g. deformed wing virus, that affects honeybees but are transmitted by mites [8]). Some iflaviruses have also been identified in ticks, for example in the tick *Hyalomma asiaticum* and in pools of different tick species [3]. Recently, a complete genome of an iflavirus has also been sequenced in an endemic Australian species, *Ixodes holocyclus* [9], in a tick species associated with marine birds, *Ixodes uriae* [10], and in the ISE6 cell line of *Ixodes scapularis* [11]. These findings illustrate two different strategies to isolate microorganisms, which each have their specific potential and interests: collecting and sequencing large pools of wild individuals (allowing to obtain sequences even for viruses that have a potentially low prevalence, as in [5] and [9]), or investigating well-characterized cell lines (as in [11]). Tick cell lines are an important tool for characterizing or experimenting on the interactions between microorganisms and mammalian systems [12] and have been screened for the presence of microorganisms, including viruses [13,14].

In this study, we took advantage of a large number of transcriptome sequencings for different tick species, either produced by our group or available in public repositories, to search for candidate sequences of *Iflaviridae.* We identified nine new genomes of iflaviruses in different tick species (or new variants constituting a quasi-species, for *Ixodes ricinus),* and discuss their evolution and phylogenetic relationships.

## 2. Materials and Methods

### 2.1 Search of Iflavirus sequences

We first searched sequences matching iflaviruses in all available public transcriptome assemblies for ticks (available from TSA, Genbank). We searched assemblies (TSA files) rather than raw read (SRA database) because i) an exhaustive search of raw reads would have implied re-assembling de novo all RNA-Seq data sets for all tick species, which represent a considerable volume of data ii) most RNA-Seq projects for ticks have been assembled *de novo* and the assemblies are available in TSA iii) virus sequences are not filtered out from *de novo* assemblies by Genbank, in our experience iv) searches of unfiltered *de novo* assemblies using raw reads, which our group produced recently for 27 different tick species [15] gave the same results as a search to TSA assemblies, regarding positive or negative detection of iflaviruses. Overall, this suggests an absence of filtering of virus sequences from the public transcriptome assemblies, and that the strategy we used was efficient to detect iflaviruses.

For each transcriptome assembly, we searched matches by using the iflavirus described from a cell line of *I. scapularis* (ISIV genome), and doing a tblastn between the poly-protein sequence (complete genome) as a query and RNA-Seq contigs as targets. This search was started with *I. ricinus*, for which a large amount of transcriptomic resources has been generated (with diverse strains and origins of samples), and then extended to other species of ticks. Some of the published assemblies for different species of the genus *Ixodes* had been produced by our research group for a phylogenetic study of hard ticks (Charrier et al., 2019). Our group has additionally recently produced three new transcriptomes for *I. ricinus* (Table 1), with specific objectives (transcriptomic study of the synganglion, transcriptome of whole nymphs, and expression profiling of different organs during feeding). These three transcriptomes have not yet been published as the three studies still need to be finalized, and will be described in upcoming manuscripts, but we used them to complete the search of iflavirus genomes with the same methods.

**Table 1.**
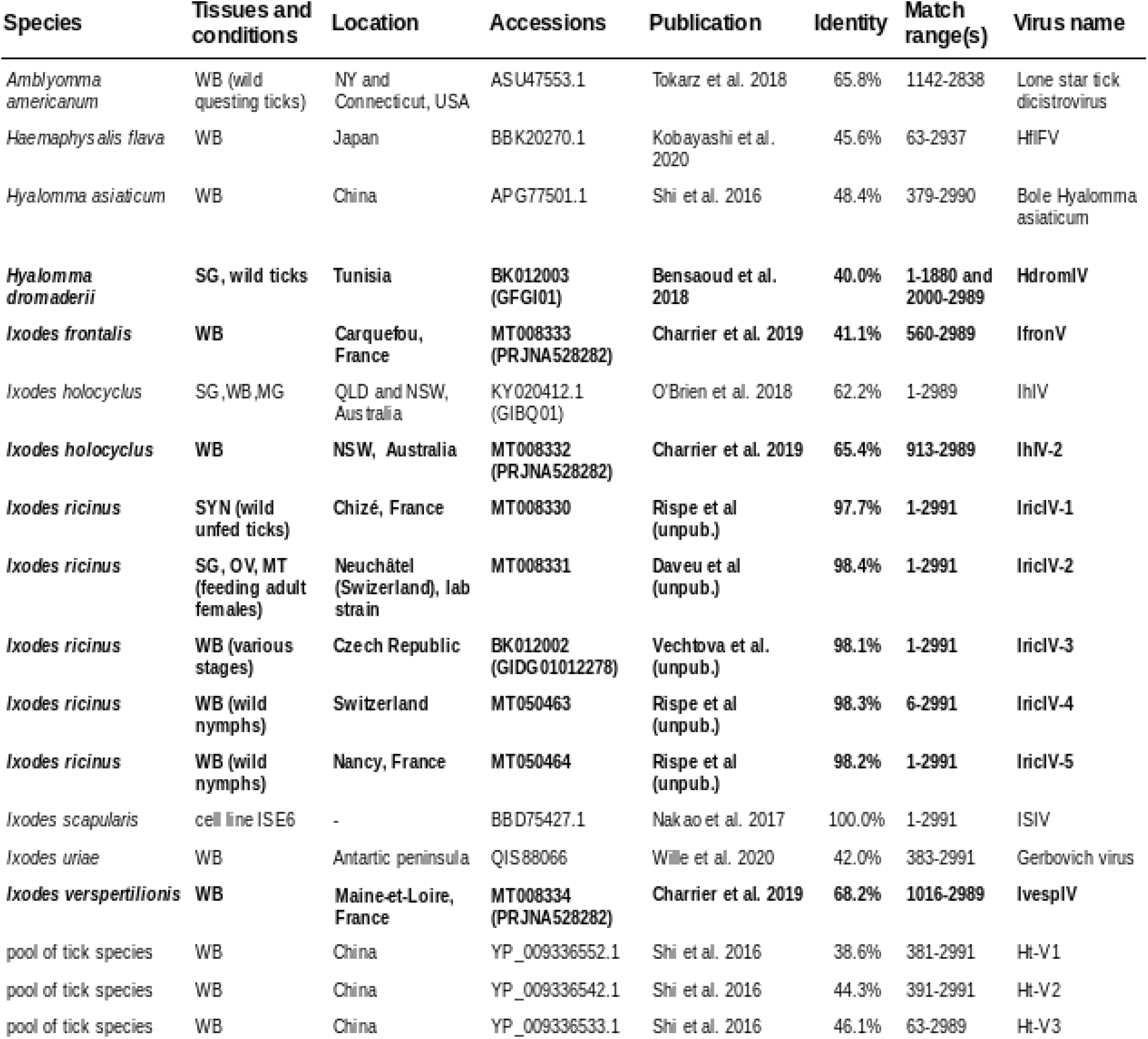
List of iflavirus sequences found in tick transcriptomes (with tblastn or blastp, using the ISIV genome, associated with the ISE6 cell line of *Ixodes scapularis* as a query). Columns: Species, Tissues (HEM: haematocytes, MG: midgut, MT: Malpighian tubules, OV: ovaries, SG: salivary glands, SYN: synganglion, WB: whole bodies) followed by details on the stages or conditions between parentheses, when available, Location of the sampling (or source of the strains), Accessions: Genbank accession and if available, related TSA or BioProject accession between parentheses, Publication (or authors of the sequences), Percent identity with ISIV -% id at the amino acid level of the first hsp (tblastn)- and range of the match, which indicates genome completion. Lines in bold correspond to the nine iflavirus genome sequences newly discovered in the present study.

### 2.2 Phylogeny methods

The phylogenetic analysis included all of the iflavirus genome sequences associated with ticks, as identified above. Based on the recent phylogenetic study of iflaviruses [16], we identified two relevant outgroups, which were also included. The first was an iflavirus associated with *Tetranychus truncatus* (representing the closest outgroup to tick iflaviruses) while the second was associated with *Apis mellifera* (Deformed wing virus, or Dfw) – this was a more distant outgroup, but still pertaining to the same cluster (Cluster 1) in the phylogeny of Kobayashi et al. [16]. The alignment of all protein sequences was done with MUSCLE in MegaX [17]. This alignment was filtered with Gblocks [18], which excluded poorly aligned regions and gaps. A phylogenetic ML tree was obtained with IQ-TREE [19]. The best model of substitution was determined with Model Finder [20], then we assessed branch support with 1000 ultrafast bootstrap replication [21]. A graphical edition of the consensus ML tree was performed with ITOL [22].

### 2.3 Test of congruence between host and virus phylogenies

To test the congruence between virus and host phylogenies, we used the cophylogeny testing tool Jane 4 [23]. As three of the iflavirus genomes were obtained from pools of tick hosts, from different genera, we could not include these three genomes and made this analysis with a subset of the tree, containg all other sequences. We also collapsed virus sequences that were nearly identical in the virus phylogeny, since they can be considered to be variants and belong to the same species. This concerned two iflaviruses associated with *I. holocyclus*, but also a group of six sequences, five being found in *I. ricinus* and the latter in a cell line of *I. scapularis* (see Results): the name given to this group was IricIV-ISIV.

### 2.4 Estimation of genetic distances and evolutionary rates

We estimated genetic distances and evolutionary rates within two ensembles of sequences (respectively ISIV + *I. ricinus* variants, and two variants associated to *I. holocyclus*) that grouped closely in the phylogenetic analyses. We first estimated genetic distances (number of substitutions per site) both at the nucleotide level and at the amino acid level, using the Maximum Composite Likelihood model [24] and the Poisson correction model [25] respectively, with the complete deletion option (all gaps excluded). This analysis was performed in MEGA X [17]. Then, we estimated ratios of non-synonymous and synonymous substitutions (dN/dS) with Codeml [26]: for this we used the one-ratio model on a sub-tree comprising the ISIV sequence plus five variants found in *I. ricinus* and a pairwise estimate of dN/dS for the two variants found in *I. holocyclus*.

## 3. Results

### 3.1 Iflavirus sequences identified

Five variants of an iflavirus with a complete or near-complete genome could be found in different transcriptomes of *Ixodes ricinu*s – see Table 1 for details. One variant was found in an assembled transcriptome of synganglions from unfed wild adult ticks (Rispe et al. in prep.), the corresponding sequence being named IricIV-1. While there was read support for the corresponding contig in eight of nine libraries, an overwhelming majority (99%) of these reads came from just one library. A second complete sequence (IricIV-2) could be retrieved from a newly obtained transcriptome of different tissues (Daveu et al. in prep.), from *I. ricinus* individuals from the lab strain from the University of Neuchâtel, Switzerland. We also explored published assemblies of *I. ricinus* transcriptomes and detected a third complete iflavirus genome (IricIV-3) in the recently published GIDG data set (TSA accession from NCBI). Last, we identified two iflavirus genomes in transcriptomes of pooled whole bodies of nymphs (Rispe et al., in prep.) collected on the vegetation, respectively in Switzerland (IricIV-4) and north-east France (IricIV-5). All viral genomes obtained in *I. ricinus* were highly similar to the ISIV sequence obtained from a cell line of *I. scapularis* (~98%, amino acid identity based on tblastn matches, while higher variation was detected at the nucleotide level, as shown below). All other data sets of reads from *I. ricinus* either showed absence of matches, or in one case, small fragmented contigs (GCJO data set) which we will not further consider here. Although the ISIV sequence was found in a cell line transcriptome of *I. scapularis*, there was no match with this sequence in *de novo* assemblies obtained for three other independently sequenced transcriptomes of *I. scapularis* (two of them at least obtained from large collections of wild ticks, including a total of ~200 males and females from three locations for the GGIX01 accession number). Then, we also screened published assemblies for other tick species, as available in Genbank-TSA: this search was positive for *I. frontalis* (IfronIV) and *I. vespertilionis* (IvespIV) but in both cases the retrieved sequences were incomplete (5’ partial), and the obtained ORF comprised respectively 2437 and 1945 amino acids (representing ~81% and ~65% of the complete genome respectively). Additionally, a 5’ partial sequence was also found in a transcriptome assembly of *I. holocyclus* (2039 amino acids, IhIV-2). This sequence (and the corresponding samples and sequencing) was different from the first iflavirus genome (IhIV) identified for that tick species. We also found a matching contig sequence in an assembled transcriptome (GFGI01 TSA assembly) of the tick *Hyalomma dromaderii.* The match was in two frames suggesting a likely frameshift, probably due to sequence errors in the reads, or in the assembly. We tentatively corrected the frameshift manually (with one base deletion) and obtained a complete iflavirus genome sequence for this species. All other assemblies, including several genera of hard and soft ticks proved negative for this search. Finally, other sequences were found after a blast (blastp) to the Genbank nucleotide database (nr): this allowed to include sequences of iflaviruses associated respectively with *Amblyomma americanum, Haemaphysalis flava, Hamephysalis asiaticum, Ixodes holocyclus, Ixodes uriae,* or with pools of tick species (Table 1).

### 3.2 Phylogeny

To obtain a comprehensive phylogeny of iflavirus genome sequences associated with ticks, we integrated all the available genome sequences (as detailed in Table 1), comprising sequences already described [3,9,11,16] and the nine sequences newly identified in the present study (Fig. 1). After filtering of regions that aligned poorly and of gaps, the alignment comprised 1149 amino acid sites. The best-fit model (BIC criteria) was LG+G4. The ML tree (consensus from 1000 bootstrap trees) showed that the five variants found in *I. ricinus* and the ISIV sequence closely grouped together – see below for details on genetic distances. This was also the case of the two variants of *I. holocyclus.* Thus two groups, respectively ISIV+*I. ricinus* variants and *I. holocyclus* variants, form a ‘quasi-species’. Another iflavirus sequence, found in *I. vespertilionis*, formed a sister group with the two groups above. Two other iflavirus genomes, respectively found in *I. frontalis* and *H. dromaderii,* grouped with Ht-V1, a sequence obtained in the study of Shi et al. [3].

**Figure 1:**
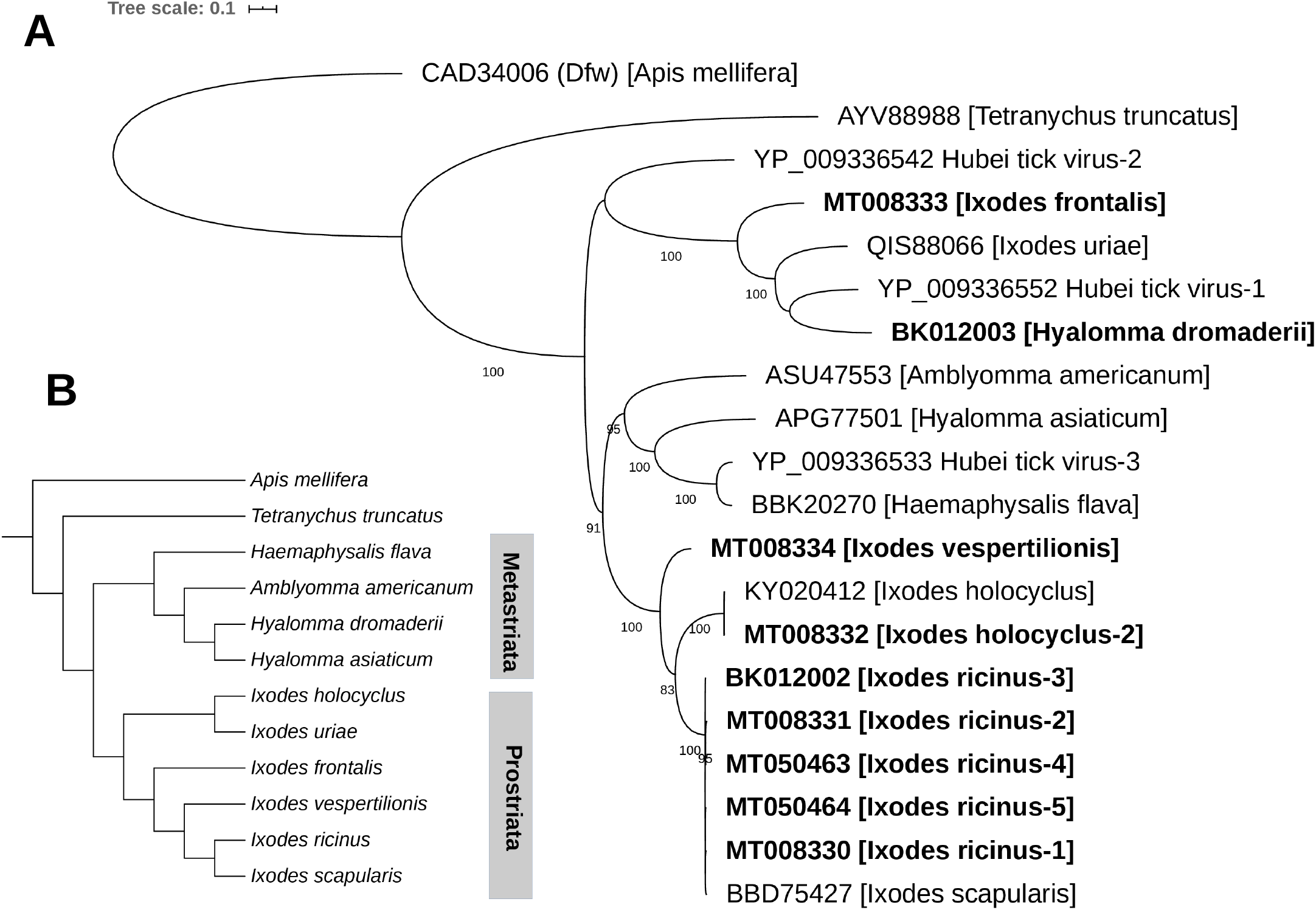
**A**, Maximum Likelihood phylogenetic tree of tick-associated iflaviruses. The tree was based on the amino acid sequence of iflavirus sequences found in the transcriptomes of different hard tick species (Acari;Parasitiform;Ixodida;Ixodidae), and rooted with two outgroups, a honeybee iflavirus (Deformed wing virus, or Dfw) and an iflavirus associated with *Tetranychus truncatus* (Acari; Acariform; Prostigmata). Host taxon is indicated between brackets. Details related to each taxon (tick host species, source of data, accession, genome completeness) are given in Table 1. Taxon names in bold correspond to the nine iflavirus sequences newly discovered in the present study. Bootstrap support is indicated at the nodes. **B**, Expected topology of the phylogenetic tree of arthropod hosts of iflaviruses included in this study, based on [14] and on the delimitation of the two groups recognized within Ixodidae (Prostriata and Metastriata).

The latter sequence was obtained from a pool of tick species which contained several species of the Metastriata (non-*Ixodes* hard ticks) or soft ticks species (Argasidae), but no species of the genus *Ixodes.* Of note, a sequence found in *A. americanum* in the study of Tokarz et al. [5] corresponds to an iflavirus although it was named “dicistrovirus” in this manuscript.

### 3.3 Test of cophylogeny

The test of cophylogeny with Jane 4 allowed us to retrieve several possible coevolutionary scenarios, which all had the same scoring and differed very little in structure (we represent one of them in Fig. 2). For all scenarios, there was an initial cophylogeny, due to the fact that all iflavirus genomes associated with ticks form a monophyletic clade. But after this ancestral node (folllowing the association of iflaviruses to ticks), all scenarios found several events of incongruence, with four instances of host switch. An additional anomaly was a “failure to diverge” between viruses associated with respectively *I. ricinus* and *I. scapularis* (see Discussion).

**Figure 2:**
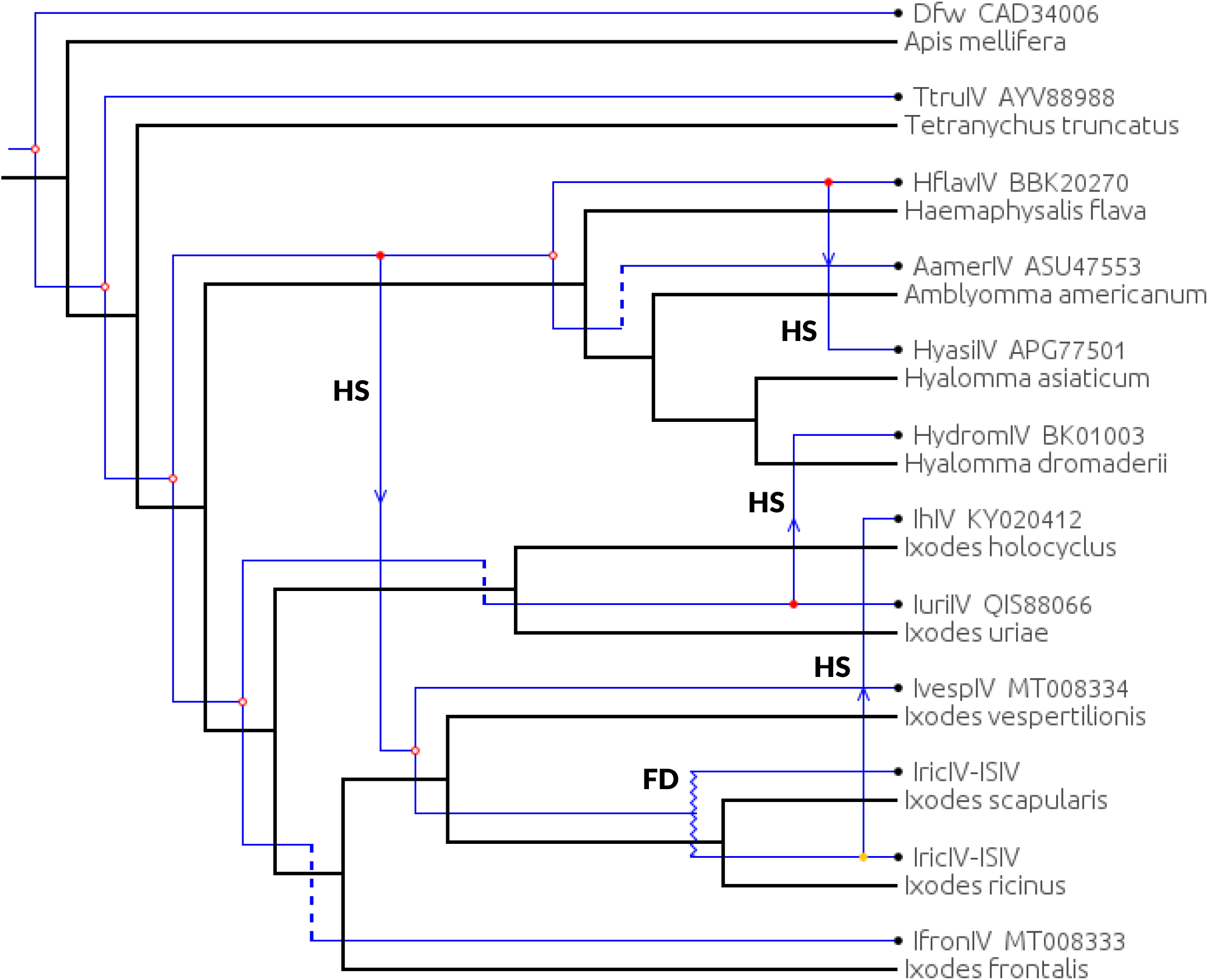
Test of the congruence between the phylogenies of tick-associated iflaviruses and of tick hosts, with Jane 4. This tree contains only part of the sequences analized in Fig. 1 (three virus genomes found in pools of tick species could not be included). Iric-ISIV represents iflavirus sequences found either in *I. ricinus* or in a cell line of *I. scapularis* (these sequences being nearly identical). Open circles at the nodes indicate co-phylogeny, HS indicates a host-shift, FD a failure to diverge.

### 3.4 Distances among iflavirus genome sequences and rates

For the group comprising the variants found in *Ixodes ricinus* plus the ISIV sequence, the estimated pairwise distances ranged between 0.010 and 0.078 at the nucleotide level, suggesting that these sequences are closely related but not identical, whereas amino acid distances were lower, ranging between 0.004 and 0.024 (Table 2). Consistently, the estimated non-synonymous to synonymous ratio was low and well below one (with the one ratio model, dN/dS=0.024). The sequences of the two variants found in *Ixodes holocyclus* were extremely close but not identical (105 differences over 6117 nucleotide positions, but only 5 differences at the amino acid level). For this pair of variants, the estimated pairwise distances were respectively 0.018 (nucleotide level) and 0.002 (amino acid level) whereas the estimated ratio of non-synonymous to synonymous ratio was also very low (paiwise ratio, dN/dS=0.014).

**Table 2.**
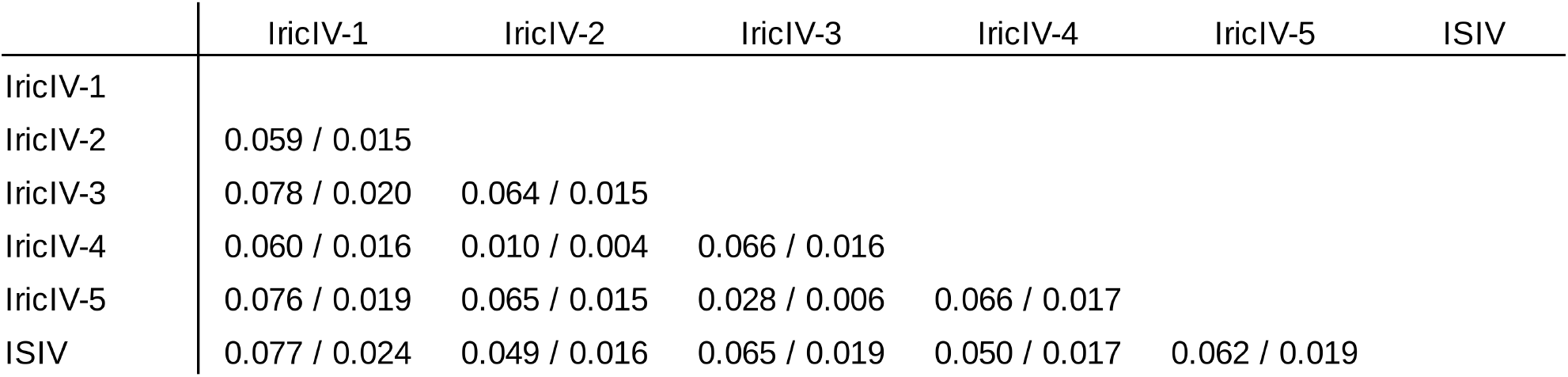
Pairwise distances among complete genomes of iflavirus associated with *Ixodes ricinus* (five variants, IricIV-1 to 5, first identified in the present study) or the *I. scapularis* cell line ISE6 (ISIV). Number of base substitutions per site (Maximum composite likelihood model, 8976 positions) followed by number of amino acid substitutions per site (Poisson correction model, 2991 positions). Analyses were conducted with MegaX.

|  | IricIV-1 | IricIV-2 | IricIV-3 | IricIV-4 | IricIV-5 | ISIV |
| --- | --- | --- | --- | --- | --- | --- |
| IricIV-1 |  |  |  |  |  |  |
| IricIV-2 | 0.059 / 0.015 |  |  |  |  |  |
| IricIV-3 | 0.078 / 0.020 | 0.064 / 0.015 |  |  |  |  |
| IricIV-4 | 0.060 / 0.016 | 0.010 / 0.004 | 0.066 / 0.016 |  |  |  |
| IricIV-5 | 0.076 / 0.019 | 0.065 / 0.015 | 0.028 / 0.006 | 0.066 / 0.017 |  |  |
| ISIV | 0.077 / 0.024 | 0.049 / 0.016 | 0.065 / 0.019 | 0.050 / 0.017 | 0.062 / 0.019 |  |

## 4. Discussion

Communities of microorganisms associated with ticks are attracting a special interest in the context of the growing concern for tick-transmitted diseases [27]. In recent years, there has been a particular interest for viruses associated with ticks, and in particular for the iflavirus family since several tick species appeared to harbour iflavirus-like viral genomes [3,5,9,11,16]. Our study, based on the exploration of assembled transcriptomes, allowed to discover nine new iflavirus genome sequences in ticks, and to perform a phylogenetic study that includes both these new genome sequences and previously published sequences. Our phylogenetic analysis showed that the iflavirus genome sequences associated to ticks closely grouped together and formed a monophyletic clade. This suggests an ancient association between this virus sub-group and ticks, which could mean relatively infrequent host-switches of the virus between major groups of arthropods. Of note however, two iflavirus-like genome sequences have been recently described for the *Antricola* genus [28], a genus of neotropical soft ticks which have a peculiar biology, being associated with hot bat caves and partially feeding on bat guano instead of an exclusive vertebrate blood diet. These two sequences (not included in our phylogeny) grouped closely with insect iflaviruses and not with sequences associated with ticks, showing that there has been more than one infection of ticks by this family of viruses. Within ticks, the virus phylogeny and the tick phylogeny were not congruent, i.e. there was no strict co-cladogenesis (Fig. 1 and Fig. 2). On the one hand, a monophyletic group comprised sequences found in *I. ricinus* (or *I. scapularis* – see below), *I. holocyclus* and *I. vespertilionis*, suggesting that they form an *“Ixodes”* subclade within iflavirus associated to ticks. However, the virus sequence found in *I. frontalis* did not group with this ensemble, and was closer to iflavirus genomes found in ticks from other genera. Also, the closer grouping of *I. holocyclus* and *I. ricinus* was inconsistent with the phylogeny of this genus [15,29], since *I. holocyclus,* a species belonging to the Australasian subgroup of *Ixodes* is phylogenetically distant from *I. ricinus* and *I. vespertilionis*. Although the data for iflaviruses in genera other than *Ixodes* is still incomplete or imprecise (e.g. several sequences came from assembled transcriptomes based on pooled species of ticks), we finally note that the sequences associated with two species of the genus *Hyalomma* did not group together. In addition the formal test of cophylogeny (Fig. 2) on a subset of virus sequences (some viruses could not be included in the test, since they could not be assigned to a precise host) suggests several events of switching of the virus between ticks species or genera over evolutionary time. A more precise picture of this dynamics will require a denser data set including more iflavirus genomes, and a larger sample of tick species.

RNA viruses have relatively fast evolving sequences, which was also observed in our data set, as we compared the virus genomes associated with different tick species. There was however a striking exception, concerning the sequence found in a cell line of *I. scapularis* (ISIV), which formed a tight cluster in the ML phylogenetic tree with five different variants identified in *Ixodes ricinus*. In fact these sequences were nearly identical at the amino acid level (>98% pairwise identity of protein sequences). Therefore, ISIV and the *I. ricinus* iflavirus sequences form together a ‘quasi-species’, a puzzling result given that the tick host species *I. scapularis* and *I. ricinus* have evolved on two different continents (respectively north-east America and Europe) and are separated by several million years of divergence. The geographic distance and the scarcity of animal (birds would be the only shared potential tick hosts) movements between Western Europe and North America leaves little chance of a recent natural inter-species contamination through shared tick hosts. We thus consider a different scenario to explain this anomaly: ISIV was found in the transcriptome of a cell line of *I. scapularis* (ISE6), but we failed to detect it in any other assembled transcriptome of this species, including a data set comprising as many as 200 adult ticks from several locations [30]. On top of that, another study assessed the presence of viral sequences in *I. scapularis* after surveying several pools of wild samples (a total of >1100 individuals, combining nymphs and adults) from different locations – New York and Connecticut, USA – but did not report any iflavirus-like sequence [5]. Of note, with the same methods, this study did identify an iflavirus genome in *Amblyomma americanum* (although it was named “dicistrovirus”, this sequence groups with iflaviridae). By contrast, the five *I. ricinus* variants found in the present study were obtained for several independent samples of wild ticks (or from one lab strain initially derived from wild ticks), which strongly supports the fact that the *I. ricinus* iflavirus variants newly discovered in our study correspond to strains of iflavirus that naturally infect this species. Overall, this apparent absence of the virus (or at least of an ISIV-like sequence) in wild individuals of *I. scapularis* combined with near-identity between ISIV and the sequences associated with *I. ricinus* could be better explained by a contamination of the *I. scapularis* ISE6 cell line with a virus associated in the field with *I. ricinus.* Such contamination could have been mediated by the establishment or maintenance of cell lines of the two species in a same lab, a possibility previously discussed for other viruses and cell lines of *I. scapularis* in the study of Alberdi et al. [13].

Much remains to know on the evolutionary dynamics of the association between iflaviruses and tick species. This first concerns the modes of transmission of the virus, and the possibility of transmission among ticks of different species (or genera) that could occur through co-feeding on the same host. An important missing piece of information in the understanding of this system is also the biological effect(s) of iflaviruses on their tick hosts, as there has not been report, to our knowledge, of symptoms in ticks infected by the virus. An element to evaluate the short-term aspects of the evolution of the virus is the level of intra-specific variability (variability between variants of the viral genome found in one tick species). Our study allows, for the first time, to evaluate this variability, since we could obtain and analyse a total of six variants for the group ISIV+ *I. ricinus,* and also two variants for *I. holocyclus*. The observed variation among strains within each of these groups was mostly constituted of synonymous mutations, and we found evidence of strong purifying selection acting between these variants. This means that at least for short time paces of evolution (within a host tick species), the rate of evolution of the protein sequence of this virus is low.

The extant data is still too scarce to determine the prevalence of the virus among tick species: negative results in several RNA-Seq data for *I. ricinus,* whereas other transcriptomes of this species allowed to detect five different iflavirus genomes, which suggests that the virus is not consistently present in tick populations (and perhaps even that its prevalence is low). The negative results obtained in most species (i.e. all transcriptomes of twelve species of the Metastriata and three sp. of *Ornithodoros*) should therefore not be taken as evidence of absence of this family of virus from these species. Our study, which surveyed different RNA-Seq data sets obtained with very heterogeneous sources of materials (cell lines, lab strains, wild ticks) allowed however to significantly enrich the range of tick species with a complete iflavirus genome. This comforts the scenario of an ancient association and relatively widespread presence of iflavirus genomes in tick species. We suggest that a more systematic use of RNA-Seq, based on large pools of wild individuals, would maximize chances of detecting that type of sequences even in case of low prevalence – see for example [31]. More generally, this would enhance progress in the description of the virome associated with ticks, comprising viruses pathogenic for their vertebrate hosts, including humans [32].

## Supporting information

Supplemental material - iflavirus sequences

## Supplemental material

the file suppl_rispe.txt contains the Genbank format sequences of the newly identified iflavirus genomes.

## Author Contributions

Conceptualization and Methodology: C.R. and O.P.; Ressources and Funding acquisition: C.H., D.S., A.J., J.-M.A., K.L., O.P.; Formal analysis and Data curation: C.R, R.D., L.S.; Writing: C.R., O.P. and R.D. All authors have read and agreed to this version of the manuscript.

## Acknowledgements

we thank O. Rais, M. Voordouw and Franck Boué who provided part of the material used for transcriptome sequencing of *I. ricinus*. We acknowledge the support of the Genotoul bioinformatics platform Toulouse Midi-Pyrénées (Bioinfo Genotoul) for bioinformatic analyses. This work was supported by the Genoscope, the Commissariat à l’Energie Atomique et aux Energies Alternatives (CEA) and France Génomique (ANR-10-INBS-09-08), by the Italian Ministry of Education, University and Research (MIUR): Dipartimenti di Eccellenza Program (2018-2022) – Dept. of Biology and Biotechnology “L. Spallanzani”, University of Pavia (to D.S.), by a grant form the Institut Carnot France Futur Elevage (XENOBIOTICK project). Thanks to Sara Moutailler (ANSES) for helpful comments on the manuscript.

